# Comparison of Generally Recognized as Safe Organic Acids for Disinfecting Fresh-cut Lettuce

**DOI:** 10.1101/348045

**Authors:** Jiayi Wang, Dongbing Tao, Yubo Liu, Siwen Han, Fenge Zheng, Zhaoxia Wu

## Abstract

In this study, we aimed to determine the organic acids (acetic, lactic, citric, malic, propionic, succinic, and tartaric acids; 1% and 0.5%, w/w or v/v) that were most effective for fresh-cut lettuce disinfection based on analysis of quality (i.e., color, electrolyte leakage, and sensory quality) and microbial examination. The results showed that these acids did not negatively affect the color quality (i.e., L^*^, a^*^, b^*^, whiteness index, and sensory color). Additionally, 0.5% lactic acid led to the lowest electrolyte leakage (0.83%), which was not significantly different (*p* > 0.05) from that of distilled water (0.46%). Lactic acid (1%) did not affect the sensory quality and led to the highest microbial reduction (1.45 log reduction in aerobic plate counts [APCs]; 2.31 log reduction in molds and yeasts [M&Y]) and was therefore recommended as the primary choice for lettuce disinfection. Malic acid (0.5%), with a 1.07% electrolyte leakage rate, 0.73 log reduction in APCs, and 1.40 log reduction in M&Y, was better than the other six acids (0.5%) and was recommended as a pH regulator and a potential synergistic agent for oxidizing sanitizers. Acetic acid (1%) negatively affected the sensory quality and led to the highest electrolyte leakage (2.90%). Microbial analysis showed that propionic acid (0.5% and 1%) was ineffective for disinfection of lettuce (*p* > 0.05); thus, acetic and propionic acids were not recommended. Our results provide insights into the choice of sanitizers and formula design in food safety.

**IMPORTANCE:** Since chlorine is forbidden in several countries, generally recognized as safe organic acids are used in minimal processing industries and in household sanitizers. The disinfection efficacy of organic acids has been studied when used alone or with oxidizing sanitizers. However, since different antibacterial mechanisms, contact time, fresh produce, and concentration have been reported, the acids most effective for single fresh produce disinfection, especially that of lettuce, an important salad vegetable, are not known. Moreover, in developing countries, because of imperfections in field management, cold chain transportation, and minimal processing industry development, the demand for low-cost household sanitizers is greater than that for minimally processed fresh produce. In this work, microbial load in lettuce was determined after disinfecting with seven GRAS organic acids. The changes in quality were also determined. These results provide insights into the choice of minimal processing sanitizers and a formula design for household sanitizers.

## 1. Introduction

Consumption of fresh produce is an important part of the daily diet, providing necessary vitamins, minerals, and cellulose. The United States Department of Agriculture recommends a daily intake of 3–5 different vegetables and 2–4 different fruits. Although fresh-cut produce offers diversification, essential nutrients, and convenience, numerous foodborne diseases related to microbial contamination (typically *Escherichia coli* O157:H7 and *Salmonella* spp.) have been reported in these products (Berger et al., 2010; Castro-Ibáñez, Gil, & Allende, 2017; Newell, Koopmans, Verhoef, Duizer, Aidara-Kane, Sprong et al., 2010; Yaron & Römling, 2014). In addition, the decay of fruits and vegetables caused by fungi is responsible for huge economic losses to growers, tradesmen, and consumers.

Chemical sanitizers have been widely used in minimal processing industries owing to their advantages of efficacy and low cost (Meireles, Giaouris, & Simões, 2016), such as chlorine, organic acid, ozone, hydrogen peroxide, sodium hypochlorite, and quaternary ammonium compounds (Castro-Ibáñez, Gil, & Allende, 2017; Ma, Zhang, Bhandari, & Gao, 2017; Ramos, Miller, Brandão, Teixeira, & Silva, 2013). However, chlorine-based agents are ineffective against some microorganisms and can lead to the generation of carcinogenic compounds (Glowacz, Colgan, & Rees, 2015). Therefore, application of nonchlorine sanitizers has attracted much attention. Several comparison studies between chlorine and other chemical sanitizers have been performed recently (Bermúdez-Aguirre & Barbosa-Cánovas, 2013; Finten, Agüero, & Jagus, 2017; Neal et al., 2012; Poimenidou et al., 2016). Among these sanitizers, most organic acids have been given the classification of “generally recognized as safe” (GRAS) by the US Food and Drug Administration and have also been used as flavoring agents and pH regulators in the food industry. Organic acids are better than sodium hypochlorite since toxic and carcinogenic compounds are not generated (Lianou, Koutsoumanis, & Sofos, 2012). Acetic, lactic, and citric acids are commonly used for the treatment of fresh-cut produce (Ölmez & Kretzschmar, 2009). The use of other acids (i.e., malic acid, propionic acid, succinic acid, and tartaric acid) as sanitizers, listed as “direct food substances affirmed as GRAS”, has also been reported in recent years (Finten, Agüero, & Jagus, 2017; Meireles, Giaouris, & Simões, 2016; Park, Baek, Kim, Kim, & Koo, 2013; Ramos-Villarroel, Martin-Belloso, & Soliva-Fortuny, 2015; Samara & Koutsoumanis, 2009). The antibacterial mechanisms of action of these compounds are based on reduced environmental and cellular pH values and increased anion accumulation (Carpenter & Broadbent, 2009; Parish, et al., 2003). Although many studies have shown good disinfection effects using different organic acids, it is unclear which acids are better since these studies used different contact times, concentrations, and tissues. Lianou, Koutsoumanis, and Sofos (2012) also believed that the decontamination efficacy of organic acids cannot be evaluated when different types of acids are used and different tissues are treated. Moreover, the use of high concentrations of organic acids, such as 5% acetic acid (Chang & Fang, 2007; Wu, Doyle, Beucgat, Wells, Mintz, & Swaminathan, 2000) and lactic acid (> 1.6%) (Van Haute, Uyttendaele, & Sampers, 2013), to achieve good bactericidal effects can yield poor organoleptics (Ölmez & Kretzschmar, 2009). In addition to microbial reduction, the effects of organic acids on physical qualities, such as color and electrolyte leakage, should also be considered. To the best of our knowledge, no reports have compared the disinfection efficacy of GRAS organic acids and considered the quality attributes described above.

Therefore, in this study, we compared the effects of seven GRAS acids (i.e., acetic, lactic, citric, malic, propionic, succinic, and tartaric acids) on microbial reduction, physical quality, and sensory quality of fresh-cut lettuce using concentrations of 0.5% and 1%. Our results provide insights into the choice of sanitizers in food safety.

## 2. Materials and Methods

### 2.1. Sample preparation

Lettuce was purchased at a local market on the day of the experiment. Two outer leaves, inner baby leaves, and the stem were removed. The remaining leaves were then rinsed in tap water for 1 min to remove the soil and cut in pieces using a square cutting edge (diameter of 5.2 cm).

Acetic acid (AA), lactic acid (LA), citric acid (CA), malic acid (MA), propionic acid (PA), succinic acid (SA), and tartaric acid (TA) were all of analytical grade and purchased from Macklin (Shanghai, China). The p*K*_a_ values of TA, LA, CA, MA, SA, AA, and PA were 2.98, 3.08, 3.14, 3.46, 3.46, 4.21, 4.75, and 4.87, respectively. The pH values of 1% TA, LA, CA, MA, SA, AA, and PA were 2.08, 2.15, 2.11, 2.22, 2.55, 2.66, and 2.78, respectively, and those for 0.5% TA, LA, CA, MA, SA, AA, and PA were 2.22, 2.36, 2.31, 2.35, 2.71, 2.81, and 2.99, respectively. The acid solutions were prepared at two concentrations (0.5% and 1.0% w/v or v/v), and the cut samples were dipped into the solutions (18 ± 1 °C) for 2 min at a ratio of 1:30 (w/v). Water was removed from the disinfected samples using a manual salad spinner, and the samples were then placed on sterile gauze for subsequent analysis.

### 2.2. Microbiological analysis

After washing, 5 g sample was evenly divided into four parts and completely immersed into an Erlenmeyer flask containing 70 mL sterile 0.9% sodium chloride solution. The samples were shaken for 3 min at 260 rpm to prepare a 15-fold dilution, and 6 mL of the suspension was added to 34 mL of 0.9% sodium chloride solution to prepare a 100-fold dilution. Other serial dilutions were prepared as needed and uniformly mixed using a vortex shaker before use. For the aerobic plate counts (APCs), 1 mL of the dilution was pour-plated into plate count agar (Hopebio, Qingdao, China), and the colonies were counted after a 2-day incubation period at 37 °C. For molds and yeasts (M&Y), Rose Bengal agar (Hopebio, Qingdao, China) was used for analysis, and the colonies were counted after incubating for 5 days at 28 °C. Samples washed with distilled water were selected as the control group. Three replicates were analyzed in duplicate, and the results were expressed as log CFU/g reduction.

### 2.3. Physical quality analyses

#### 2.3.1. Color

Five leaves in each of the treatment and control (distilled water) groups were randomly selected for color analysis. CIELab color space was selected, and values for L^*^, a^*^, and b^*^ were determined for two locations on each leaf using a colorimeter (CR400; Konica Minolta, Osaka, Japan). Among the three parameters, negative to positive values for L^*^ represented darkness to lightness; values for a^*^ represented green to red; and values for b^*^ represented blue to yellow. ΔE and whiteness index (WI) were selected to evaluate overall color difference and whiteness extent after washing and were calculated using the following formula:

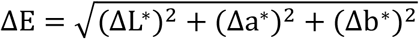

where ΔL*, Δa*, and Δb^*^ represent the differences between the treatment and control groups.

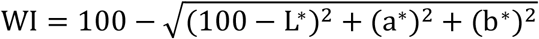

The colorimeter was calibrated using a white standard plate (*Y* = 82.80, *x* = 0.3194, *y* = 0.3264) before every use. Each experiment was independently performed three times.

#### 2.3.2. Electrolyte leakage

Electrolyte leakage was selected to estimate the extent of surface damage after washing with acid solution, as described by Gómez-López, Marín, Medina-Martínez, Gil, and Allende (2013) and Zhang and Yang (2017), with some modifications. Briefly, 5 g washed samples were immersed in 250 mL distilled water (1–2 μs) for 20 s to remove residual acid solution, which could interfere with the conductivity measurement. After centrifugation in the manual salad spinner, the samples were immersed in 150 mL distilled water, and the conductivity was measured after 0.5 and 30 min. The samples were then incubated at -20 °C for 24 h and thawed overnight at room temperature. Samples washed with distilled water were used as a control group. Each experiment was independently performed three times. Electrolyte leakage was calculated using the following formula:

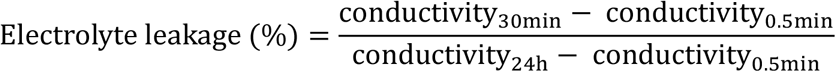

### 2.4. Sensory analysis

After washing, the samples were placed on a tray with marks at the bottom, and the order of the group was randomly reorganized. Thirteen trained members were invited to evaluate the sensory quality of the samples, including off-odor, color, texture, and overall acceptability. The scores had five rankings (Zhang & Yang, 2017), as follows. For off-odor, color, and texture, 1 indicated too many defects, 3 indicated some defects, 5 indicated moderate defects, 7 indicated few defects, and 9 indicated no defects; for overall acceptability, 1 indicated very poor, 3 indicated quite poor, 5 indicated neutral, 7 indicated quite good, and 9 indicated very good.

### 2.5. Statistical analysis

The data were compared with analysis of variance using SPSS v.20 software (SPSS Inc., Chicago, IL, USA), and the differences (*P* < 0.05) in means were analyzed by Duncan’s multiple range tests.

## 3. Results and Discussion

### 3.1. Effects of GRAS organic acids on color and electrolyte leakage

Because damage to leaf tissue caused by acid solutions must be greater than or equal to that by distilled water, the distilled water group was selected as the control group. Compared with the distilled water group, there were no significant changes (Table 1) in L*, a*, and b^*^ values; in particular, L^*^ and a*, which are two key indicators used to evaluate whether green leafy vegetables are discolored after washing with sanitizers, were not altered. The nonsignificant differences in WI values (Table 1) also indicated that these acids did not discolor the lettuce. For ΔE values, these acids did not significantly affect the overall visual quality of the lettuce (Table 1). However, organic acids, primarily AA and LA, have negative effects on the color quality of meat products (Lianou, Koutsoumanis, & Sofos, 2012), such as pork (Castelo, Kang, Siragusa, Koohmaraie, & Berry, 2001; Lin & Chuang, 2001) and beef (Pipek, Šikulová, Jeleníková, & Izumimoto, 2005), with less red than untreated samples. This appearance could be attributed to protein denaturation caused by pH reduction (González-Fandos & Dominguez, 2006). Similarly, when hydrogen peroxide and LA are used in combination, the quality characteristics of apples are not affected, but lettuce shows moderate browning (Lianou, Koutsoumanis, & Sofos, 2012; Lin, Moon, Doyle, & Mcwatters, 2002; Mcwatters, Doyle, Walker, Rimal, & Venkitanarayanan, 2002), indicating that chemical sanitizers have different effects on the visual qualities of different types of produce. The waxy layer and fluff on the surface and the more curled surface structure compared with that of meat products may be responsible for the observed color retention after treatment with organic acids. Moreover, sanitizers with oxidizing ability are expected to exhibit stronger decolorizing effects than organic acids. In a study by Martín-Diana, Rico, Barry-Ryan, Frías, Henehan, and Barat (2007), the -a^*^ values of fresh-cut lettuce were found to be decreased after treatment with 120 ppm chlorine for 1 min. Bermúdez-Aguirre and Barbosa-Cánovas (2013) found that L^*^ and WI values increased after exposure to 15 ppm ozone, whereas 0.5–1.5% CA did not interfere with the visual quality of the produce. Poimenidou et al. (2016) reported that the b^*^ values of lettuce, after storage for 6 days, were 37.1, 22.5, and 24.1 following washing with 300 ppm sodium hypochlorite, 2% lactic acid, and vinegar (6% AA), respectively. These phenomena may be attributed to the direct damage of the lettuce surface by oxidizing agents, such as ozone, which causes lipid oxidation in the cell envelope (Khadre, Yousef, & Kim, 2001), unlike organic acids inducing irreversible denaturation of intracellular acid-sensitive proteins (Ricke, 2003) without inducing membrane damage (Cherrington, Hinton, Pearson, & Chopra, 1991; Thompson & Hinton, 1996).

**Table 1.** Effects of different organic acids on the physical qualities of fresh-cut lettuce

| Treatment | L<sup>*</sup> | -a<sup>*</sup> | b<sup>*</sup> | △E | WI | Electrolyte leakage (%) |
| --- | --- | --- | --- | --- | --- | --- |
| DI | 51.03±4.55<sup>ab</sup> | 19.60±0.80<sup>abcdef</sup> | 33.62±2.63<sup>abcd</sup> |  | 37.32±1.93<sup>a</sup> | 0.46±0.16<sup>a</sup> |
| 1% CA | 49.10±0.86<sup>ab</sup> | 19.20±0.46<sup>abcd</sup> | 33.41±0.60<sup>abcd</sup> | 3.85±3.47<sup>a</sup> | 36.15±1.10<sup>a</sup> | 1.13±0.25<sup>bcd</sup> |
| 0.5% CA | 51.14±2.48<sup>ab</sup> | 19.30±0.79<sup>abcd</sup> | 33.37±2.32<sup>abcd</sup> | 4.11±3.18<sup>a</sup> | 37.70±0.85<sup>a</sup> | 1.17±0.26<sup>bcd</sup> |
| 1% MA | 50.77±1.06<sup>ab</sup> | 19.51±0.76<sup>abcde</sup> | 32.43±1.71<sup>ab</sup> | 5.29±3.47<sup>a</sup> | 37.88±0.30<sup>a</sup> | 1.28±0.08<sup>bcd</sup> |
| 0.5% MA | 49.32±1.64<sup>ab</sup> | 19.16±0.85<sup>abcd</sup> | 31.92±2.12<sup>a</sup> | 4.62±4.97<sup>a</sup> | 37.08±1.13<sup>a</sup> | 1.07±0.20<sup>bc</sup> |
| 1% LA | 48.18±1.69<sup>a</sup> | 18.57±0.32<sup>a</sup> | 31.53±0.31<sup>a</sup> | 3.87±3.66<sup>a</sup> | 36.55±1.32<sup>a</sup> | 1.24±0.10<sup>bcd</sup> |
| 0.5% LA | 50.99±2.47<sup>ab</sup> | 19.08±0.79<sup>ab</sup> | 33.32±1.31<sup>abcd</sup> | 3.99±2.68<sup>a</sup> | 37.72±2.02<sup>a</sup> | 0.83±0.19<sup>ab</sup> |
| 1% TA | 49.67±1.66<sup>ab</sup> | 19.14±0.33<sup>acd</sup> | 32.40±0.76<sup>ab</sup> | 4.61±4.13<sup>a</sup> | 37.15±0.95<sup>a</sup> | 1.31±0.16<sup>cde</sup> |
| 0.5% TA | 50.62±1.90<sup>ab</sup> | 19.45±0.38<sup>abcde</sup> | 32.75±0.47<sup>abc</sup> | 5.11±3.49<sup>a</sup> | 37.62±1.17<sup>a</sup> | 1.34±0.26<sup>cde</sup> |
| 1% SA | 50.68±2.18<sup>ab</sup> | 19.93±0.64<sup>bcdef</sup> | 33.36±1.66<sup>abcd</sup> | 3.30±1.47<sup>a</sup> | 37.17±0.66<sup>a</sup> | 1.19±0.16<sup>bcd</sup> |
| 0.5% SA | 50.58±1.41<sup>ab</sup> | 19.12±0.68<sup>abc</sup> | 32.12±0.94<sup>a</sup> | 4.10±3.37<sup>a</sup> | 38.02±1.15<sup>a</sup> | 0.89±0.27<sup>bc</sup> |
| 1% PA | 52.68±2.53<sup>b</sup> | 20.55±0.38<sup>ef</sup> | 36.11±1.20<sup>d</sup> | 3.85±3.13<sup>a</sup> | 36.99±1.14<sup>a</sup> | 1.57±0.46<sup>de</sup> |
| 0.5% PA | 50.85±1.08<sup>ab</sup> | 20.65±0.31<sup>f</sup> | 35.90±0.33<sup>d</sup> | 4.63±0.36<sup>a</sup> | 35.72±0.84<sup>a</sup> | 1.59±0.30<sup>de</sup> |
| 1% AA | 50.82±3.35<sup>ab</sup> | 20.24±0.28<sup>cdef</sup> | 35.45±1.82<sup>cd</sup> | 2.57±2.03<sup>a</sup> | 36.02±1.56<sup>a</sup> | 2.90±0.29<sup>f</sup> |
| 0.5% AA | 51.36±0.28<sup>ab</sup> | 20.27±0.39<sup>def</sup> | 35.00±1.07<sup>bcd</sup> | 4.15±0.51<sup>a</sup> | 36.74±0.50<sup>a</sup> | 1.74±0.17<sup>e</sup> |

Values (means ± SDs) in the same column with different letters indicate significant differences ( $p < 0.05$ ). DI, distilled water; CA, citric acid; MA, malic acid; LA, lactic acid; TA, tartaric acid; SA, succinic acid; PA, propionic acid; AA, acetic acid.

Scanning electron microscopy (SEM) and transmission electron microscopy can be used to observe changes in surface structure and organelles, respectively. However, electron microscope technologies, even environmental SEM, are unsuitable for evaluating the extent of damage to lettuce because of the requirement for dehydration during SEM sample preparation. Therefore, electrolyte leakage was selected to evaluate the extent of damage after washing with organic aids. In our study, electrolyte leakage (Table 1) was significantly altered (*p* < 0.05) after washing with all acid solutions, except 0.5% LA. Among these acids, 1% AA led to the highest electrolyte leakage (2.91%; *p* < 0.05). However, in order to achieve sufficient disinfection, the concentration of AA used in some studies exceeded 1%; for example, 1.9% AA yielded a 2.3 log reduction in aerobic bacteria (Vijayakumar & Wolfhall, 2002), 4% AA yielded a 3.37 log reduction in mesophilic aerobic plate counts (Nascimento, Silva, Catanozi, & Silva, 2003), and 5% AA yielded a 3 log reduction in *E. coli* O157:H7 (Chang & Fang, 2007). Moreover, significant differences were only observed for the AA groups, indicating that electrolyte leakage may be accelerated as the concentration increases. Surface damage caused by chemical sanitizers is inevitable, although electrolyte leakage was not significantly altered by 0.5% LA. Electrolyte leakage values for samples treated with 1% CA, 0.5% CA, 1% MA, 0.5% MA, 1% LA, 1% SA, and 0.5% SA were 1.13%, 1.17%, 1.28%, 1.07%, 1.24%, 1.19%, and 0.89% respectively, which were not significantly different from those with 0.5% LA.

### 3.2. Comparison of disinfection efficacy

After washing with organic acids, APCs were significantly reduced (*p* < 0.05; Table 1), except for those following treatment with 0.5% TA, 0.5% PA, or 1% PA. For PA, the M&Y counts were also not significantly reduced, suggesting that PA was not ineffective for lettuce disinfection. Similarly, in another study by Fernandes, Flick, Cohen, and Thomas (1998), the counts of aerobic bacteria on catfish fillets were significantly reduced after spraying for 20 min using 2% and 4% PA, whereas 1% PA was found to be ineffective. Another study found that *Listeria monocytogenes* was simulated to grow on lettuce after washing with 0.5% PA, whereas 1% PA significantly reduced the counts of this bacterium; the authors suggested that this result may be explained by the fact that *L. monocytogenes* is more resistant to 0.5% PA than native microflora and is more competitive, whereas 1% PA can create an acidic environment that exceeds the upper limit of resistance of the bacterium (Samara & Koutsoumanis, 2009). Except for LA, our results also showed that there were no significant differences in APC log reductions between high and low concentrations of all acid solutions. This phenomenon was also found for *E. coli* O157:H7 using TA, SA, AA, LA, and MA at concentrations of 1% and 2% (Huang & Chen, 2011).

**Table 2.**
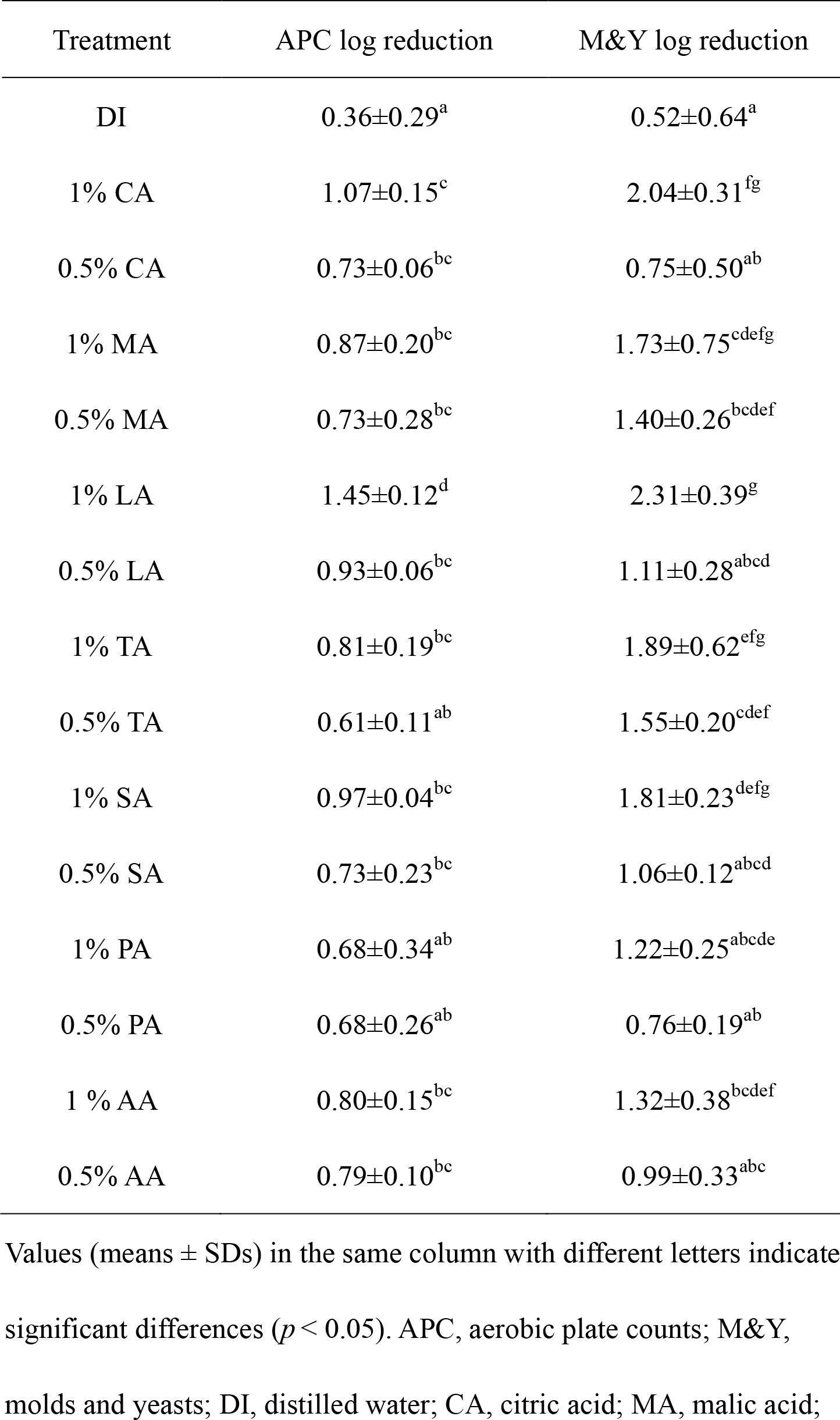
Effects of different organic acids on the natural microbiota of fresh-cut lettuce

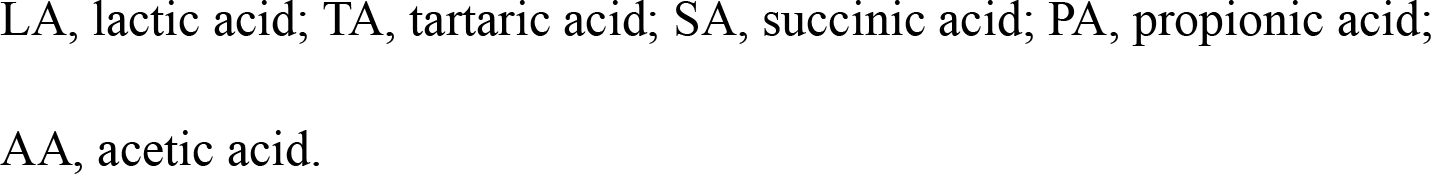

For M&Y, 0.5% CA, 0.5% LA, 0.5% SA, 0.5% AA, 0.5% PA, and 1% PA showed no advantages over distilled water. In a study by Zhang and Yang (2017), 0.5% CA was also found to be ineffective for disinfecting M&Y on lettuce. Similar to APCs, M&Y reduction was not significantly different for acids with different concentrations (0.5% and 1%), except for LA and CA. Moreover, combinations of low-concentration organic acids and oxidizing sanitizers were better than the use of a single compound alone. Interestingly, 0.6% CA and 2% H_2_O_2_ can reduce AMC and M&Y by less than 1 log when used alone, but by 2.26 and 1.28 log, respectively, when used in combination (Zhang & Yang, 2017). Moreover, combined use of chlorine and electrolyzed water with low-concentration organic acids also exhibits good disinfection effects (Huang, Hung, Hsu, Huang, & Hwang, 2008; Inatsu, Bari, Kawasaki, Isshiki, & Kawamoto, 2005). According to our results, among the low-concentration groups, only 0.5% TA and 0.5% MA significantly reduced M&Y (*p* < 0.05), and 0.5% TA did not significantly reduce APCs. Therefore, 0.5% MA was recommended as a pH regulator and synergistic agent for oxidizing sanitizers. Additionally, our results showed that M&Y reduction was better than APC reduction, consistent with a previous study (Nascimento, Silva, Catanozi, & Silva, 2003). Organic acids are also better than several oxidizing sanitizers, such as ozone and chlorine, for disinfection of M&Y. Wei, Zhou, Zhou, and Gong (2007) found that 20 mg/L ozonated water reduced M&Y on strawberries by 0.78 log and that 10 mg/L ozonated water reduced M&Y on lettuce by 0.99 log. On durum wheat, washing with 16.5 mg/L ozonated water only reduced M&Y by 0.5 log (Dhillon, Wiesenborn, Wolf-Hall, & Manthey, 2009). Martínez-Sánchez, Allende, Bennett, Ferreres, and Gil (2006) compared the disinfection efficacies of chlorine, aqueous ozone, and Purac (2% LA), and their results showed that Purac was better than ozone and chlorine for controlling M&Y during storage.

Among these acids, 1% LA was shown to be the most effective treatment solution, with 1.45 and 2.31 log reductions in APCs and M&Y, respectively (*p* < 0.05). The dissociation constant (p*K*_a_) is a key indicator to evaluate the dissociation extent of organic acids in aqueous solution; this value is dependent on the pH value and independent of concentration. The antibacterial activities of organic acids are traditionally attributed to cellular anion accumulation, which is determined by the proportion of undissociated molecules. Compared with dissociated anions, undissociated acidic molecules have stronger lipophilicity, allowing them to penetrate the microbial cell membrane more easily. After penetration, the higher intracellular pH in the environment will promote acid molecule dissociation, and the dissociated anions will accumulate in the cell and exert toxic effects on DNA, RNA, and ATP synthesis (Cherrington, Hinton, Mead, & Chopra, 1991). However, Ricke (2003) suggested that the relationship between energy dissipation and ATP production is complex and proposed that acid-sensitive protein denaturation and changes in osmotic pressure are major antibacterial mechanisms of action. Comparison of the efficacies of different acids, however, must be made at the same concentration (mol/L; i.e., the p*K*_a_ value cannot be used to evaluate the bactericidal efficiency of organic acids under different molar concentrations) (Cherrington, Hinton, Mead, & Chopra, 1991a). However, the concentration used in minimal processing industries is often w/w or v/v, which was also used in this study. In addition to p*K*_a_ values, differences in anion structure also affect the disinfection efficacy; for example, cinnamic acid (p*K*_a_ 4.4) is more effective than benzoic acid (p*K*_a_ 4.2) (Cherrington, Hinton, Mead, & Chopra, 1991a). Moreover, not all organic acids exert their activity through anion accumulation; as an example, CA acts more as a chelator to sequester metal ions (e.g., Ca^2+^, Mg^2+^, Fe^3+^) from the external medium required for bacterial homeostasis (Brul & Coote, 1999; Lianou, Koutsoumanis, & Sofos, 2012). Therefore, the antibacterial mechanism of action of organic acid is complex, and the disinfection efficacy of 1% LA cannot be explained through a single mechanism. We suggest that these results may be due to the relatively stronger pH-lowering activity of 1% LA, the relatively lower molecular weight (i.e., easier penetration of the membrane), the synergistic effects of H+ and accumulated lactate anions, and the potential binding of targets (intracellular protein) by lactate anions. Although LA had the best disinfection efficacy, LA was not better than the other organic acids when used for disinfection of some pathogens. For example, Akbas and Olmez (2007) reported that 0.5% and 1% LA reduced *L. monocytogenes* on lettuce by 2.7 and 2.9 log, respectively, which was not significantly different from the results for AA and CA at concentrations of 0.5% and 1%. Additionally, Huang and Chen (2011) found that 1% LA reduced *E. coli* O157:H7 on spinach by 1.9 log, which was not significantly different from the results for CA, MA, TA, and AA. Ganesh et al. (2010) also found that there were no significant differences between MA and LA in the control of *Salmonella Typhimurium* on spinach. Furthermore, when using sodium chlorite and organic acids in combination, there were no significant differences in *E. coli* O157:H7 populations between AA, MA, CA, TA, LA, SA, and PA (Inatsu, Bari, Kawasaki, Isshiki, & Kawamoto, 2005). Therefore, our results suggested that LA was better than the other GRAS acids for disinfecting natural microbiota present on lettuce.

### 3.3. Sensory quality

For off-odor, 1% TA, 0.5% SA, 1% SA, and 1% AA negatively affected the organoleptic properties of the lettuce (*p* < 0.05; Table 3). Similarly, the overall acceptability scores of 0.5% SA, 1% SA, and 1% AA were all 3.92 and were significantly different (*p* < 0.05) from that (6.54) of the control group. The off-odor scores for CA, LA, and MA were not significantly reduced, consistent with previous studies (Finten, Agüero, & Jagus, 2017; Samara & Koutsoumanis, 2009). When considering organoleptic properties and bactericidal efficacy, CA was found to be better than chlorine-based sanitizers. For example, in a study by Finten, Agüero, and Jagus (2017), CA was superior to sodium hypochlorite for the control of *E. coli* and *L. innocua* during refrigerated storage, whereas organoleptic score was not significantly different from that on using sodium hypochlorite. Additionally, when used in combination, the sensory quality of the combined sanitizer (hydrogen peroxide + CA) was not significantly changed, and the combination had the best disinfection efficacy compared with other sanitizers (hydrogen peroxide and hydrogen peroxide + electrolyzed water) and distilled water (Zhang & Yang, 2017). Interestingly, the off-odor score of PA was not significantly different from that of the control group, and PA had the highest overall acceptability, with the exception of 0.5% LA, inconsistent with a study by Samara and Koutsoumanis (2009). Notably, during solution preparation, PA had a very poor odor, similar to that of vomit. This result may be attributed to the volatilization in air and the reaction between lettuce aroma and PA. Consistent with the instrument color analysis, sensory color was also not significantly changed by these seven organic acids. According to our results, AA was not better than the other acids in terms of microbial reduction and sensory quality. Moreover, AA is not as effective as other acids in disinfecting pathogens (Akbas & Olmez, 2007). However, the low cost and availability as a common household item (vinegar) are important advantages of AA.

**Table 3.** Effects of different organic acids on the sensory qualities of fresh-cut lettuce

| Treatment | Off-odor | Color | Texture | Overall liking |
| --- | --- | --- | --- | --- |
| DI | 6.69±2.29<sup>e</sup> | 6.08±2.53<sup>a</sup> | 7.00±1.83<sup>a</sup> | 6.54±2.47<sup>b</sup> |
| 1% CA | 5.46±1.85<sup>bcde</sup> | 6.69±2.14<sup>a</sup> | 6.69±2.29<sup>a</sup> | 5.46±2.03<sup>ab</sup> |
| 0.5% CA | 5.92±2.66<sup>cde</sup> | 7.00±1.41<sup>a</sup> | 7.15±1.91<sup>a</sup> | 5.62±2.22<sup>ab</sup> |
| 1% MA | 5.77±2.39<sup>cde</sup> | 6.38±1.71<sup>a</sup> | 6.38±1.71<sup>a</sup> | 5.15±1.52<sup>ab</sup> |
| 0.5% MA | 5.15±2.23<sup>abcde</sup> | 7.31±1.60<sup>a</sup> | 7.00±2.00<sup>a</sup> | 5.46±1.85<sup>ab</sup> |
| 1% LA | 5.31±2.81<sup>bcde</sup> | 6.54±1.85<sup>a</sup> | 6.69±1.60<sup>a</sup> | 5.46±2.33<sup>ab</sup> |
| 0.5% LA | 6.23±1.92<sup>de</sup> | 7.15±2.08<sup>a</sup> | 7.62±1.71<sup>a</sup> | 6.23±1.74<sup>b</sup> |
| 1% TA | 4.08±2.40<sup>abcd</sup> | 6.23±2.24<sup>a</sup> | 6.23±2.09<sup>a</sup> | 4.54±2.47<sup>ab</sup> |
| 0.5% TA | 5.15±2.23<sup>abcde</sup> | 6.38±2.06<sup>a</sup> | 6.54±1.85<sup>a</sup> | 5.15±2.51<sup>ab</sup> |
| 1% SA | 3.00±2.58<sup>a</sup> | 6.54±1.66<sup>a</sup> | 6.54±2.03<sup>a</sup> | 3.92±2.66<sup>a</sup> |
| 0.5% SA | 3.77±2.65<sup>abc</sup> | 7.15±1.52<sup>a</sup> | 6.23±1.54<sup>a</sup> | 3.92±2.10<sup>a</sup> |
| 1% PA | 5.46±2.03<sup>bcde</sup> | 7.62±1.50<sup>a</sup> | 7.00±1.63<sup>a</sup> | 5.92±2.10<sup>ab</sup> |
| 0.5% PA | 5.46±2.96<sup>bcde</sup> | 6.85±1.52<sup>a</sup> | 6.69±1.80<sup>a</sup> | 5.62±2.50<sup>ab</sup> |
| 1 % AA | 3.46±2.03<sup>ab</sup> | 6.38±1.89<sup>a</sup> | 6.38±1.89<sup>a</sup> | 3.92±2.40<sup>a</sup> |
| 0.5% AA | 5.00±2.94<sup>abcde</sup> | 6.23±2.09<sup>a</sup> | 7.15±1.91<sup>a</sup> | 5.31±1.80<sup>ab</sup> |

Values (means ± SDs) in the same column with different letters indicate significant differences ( $p < 0.05$ ). DI, distilled water; CA, citric acid; MA, malic acid; LA, lactic acid; TA, tartaric acid; SA, succinic acid; PA, propionic acid; AA, acetic acid

In conclusion, 1% LA was recommended as a primary choice for lettuce disinfection. Among the low-concentration groups, 0.5% MA was recommended as a pH regulator or synergistic agent for oxidizing sanitizers. Moreover, synergistic effects were observed between different acids, such as AA and LA. However, our study only compared the disinfection efficacies of different acids when used alone. Future studies are needed to examine the synergistic effects among LA, CA, and MA. The antibacterial mechanism of action of organic acids is complex and has not been fully elucidated. With the development of omics technologies (e.g., genomic, transcriptomic, proteomic, and metabolomic), the mechanisms should be further clarified at the molecular level. Additionally, metagenomics could be used to determine which types of microbes are eliminated in order to design more effective sanitizer formulas.

## Acknowledgments

We would like to thank Xiaofei Yang, Hanling Liang, Weijia Zhang, Shan Wang, Chen Li, Wanning Zhao, Chen Yang, Yu Fu, Yubo Liu, Tian Zhang, Tianyu Wang, and Siwen Han for assistance during sensory analysis.

